# β-Amyloid and Glutathione Dysregulation Cooperatively Drive Lipid Peroxidation and Ferroptosis in Neuron-Like Cells

**DOI:** 10.64898/2026.04.15.718809

**Authors:** Kazi Rafsan Radeen, Caili Hao, Zongbo Wei, Xingjun Fan

**Affiliations:** Department of Cellular Biology and Anatomy, Medical College of Georgia at Augusta University, Augusta, Georgia, USA

## Abstract

Alzheimer’s disease (AD) is a progressive neurodegenerative disorder characterized by β-amyloid (Aβ) accumulation and oxidative stress, with aging being its greatest risk factor. Age-related decline in antioxidant defenses, particularly glutathione (GSH), may increase neuronal vulnerability to Aβ-mediated toxicity; however, the mechanisms linking redox dysregulation to neuronal death remain incompletely understood. In this study, we investigated how impaired GSH homeostasis influences neuronal susceptibility to Aβ-associated injury. Human SH-SY5Y neuron-like cells were engineered to express either wild-type APP_695_ or the familial AD-associated APP_Swe/Ind_ mutant, and intracellular GSH depletion was induced using both pharmacological and genetic approaches. GSH depletion markedly sensitized APP_Swe/Ind_-expressing cells to cell death, accompanied by increased plasma membrane lipid peroxidation, elevated malondialdehyde (MDA) and 4-hydroxynonenal (4-HNE) levels, and enhanced lactate dehydrogenase (LDH) release. This cell death was not prevented by the pan-caspase inhibitor Z-VAD-FMK but was effectively rescued by the ferroptosis inhibitors ferrostatin-1 (Fer-1) and liproxstatin-1 (Lip-1), indicating a ferroptotic mechanism. Similar ferroptotic responses were observed when Aβ oligomers were combined with intracellular GSH depletion. Mechanistically, Aβ and GSH depletion synergistically increased transferrin receptor-1 expression and intracellular iron levels while markedly suppressing glutathione peroxidase 4 (GPX4), a central regulator of ferroptosis. Importantly, inhibition of autophagy with bafilomycin A1 restored GPX4 expression and rescued cells from ferroptotic death, suggesting that autophagy-mediated GPX4 degradation contributes to this process. Collectively, our findings demonstrate that GSH dysregulation synergizes with Aβ to induce lipid peroxidation and ferroptosis in neuron-like cells. These results identify impaired redox homeostasis as a critical driver of neuronal vulnerability in AD and suggest that preserving GSH levels or targeting ferroptotic pathways may offer promising therapeutic strategies for neurodegeneration.

## Introduction

Alzheimer’s disease (AD) is the most common form of dementia and represents a progressive, irreversible neurodegenerative disorder characterized by the accumulation of β-amyloid (Aβ) plaques and neurofibrillary tangles [1]. Clinically, AD manifests a relentless decline in memory, executive function, and cognitive function, ultimately compromising an individual’s ability to perform daily activities. The global burden of this disease is staggering, as, by the year 2050, over a hundred million people are estimated to be afflicted with AD, placing an immense strain on healthcare systems, caregivers, and society at large [2].

Advancing Age is the single greatest risk factor for AD. The incidence of AD rises exponentially after the age of 65 [3, 4] underscoring a strong biological link between aging processes and neurodegeneration. Rather than serving merely as a chronological marker, aging drives a series of interconnected molecular and cellular alterations that create a permissive environment for AD pathology. These age-associated changes include impaired mitochondrial function, diminished DNA damage repair capacity, reduced proteostatic control, and a gradual decline in endogenous antioxidant defenses [5, 6]. Collectively, these alterations weaken neuronal resilience and increase susceptibility to neurodegenerative insults.

Among the mechanisms connecting aging to AD, oxidative stress has emerged as a central and unifying pathway [7]. The brain is uniquely vulnerable to oxidative damage because of its high oxygen consumption, abundance of polyunsaturated fatty acids (PUFA) prone to lipid peroxidation, and relatively modest antioxidant enzyme capacity compared to other tissues [8]. With advancing age, antioxidant defenses naturally decline, leading to an accumulation of oxidative damage to lipids, proteins, and DNA [7]. This progressive oxidative burden is widely believed to represent a critical mechanistic bridge linking normal aging to the onset and progression of AD [4, 5, 7, 9, 10].

Oxidative stress and Aβ pathology are closely interconnected. Aβ peptides, generated through sequential β- and γ-secretase cleavage of amyloid precursor protein (APP), particularly the aggregation-prone Aβ species, constitute a defining feature of AD [11, 12]. Although extracellular plaques are hallmarks of the disease, soluble Aβ oligomers are now recognized as the primary neurotoxic species, disrupting synaptic transmission, calcium homeostasis, and redox balance prior to overt neuronal loss [13]. Importantly, oxidative stress promotes amyloidogenic APP processing by increasing β-secretase (BACE1) activity while reducing α-secretase activity, thereby favoring Aβ generation [14, 15]. Oxidative modifications of Aβ further enhance its aggregation propensity and toxicity [9], establishing a feed-forward cycle between oxidative stress and amyloid pathology.

Central to the regulation of cellular redox balance is glutathione (GSH), the most abundant intracellular antioxidant. GSH maintains mitochondrial integrity, detoxifies reactive oxygen species (ROS), and modulates redox-sensitive signaling pathways in neurons and glia [16]. Notably, brain GSH levels decline during normal aging, reducing the capacity to buffer oxidative stress [17, 18]. This age-related decline in brain GSH has been consistently observed in both animal models and human studies [3, 19, 20]. In vivo magnetic resonance spectroscopy (¹H-MRS) have reveals age-dependent reductions in GSH in metabolically active brain regions, including the hippocampus, posterior cingulate cortex, and frontal cortex [17, 18, 21], regions that are also among the earliest and most severely affected in AD [18, 21]. These findings suggest that GSH depletion may contribute to regional vulnerability in neurodegeneration.

Despite growing recognition of GSH dysregulation in aging and AD, the precise molecular mechanisms by which impaired GSH homeostasis influences neuronal vulnerability under AD-related stress remain incompletely understood. Identifying how age-related redox imbalance intersects with amyloid pathology is critical for developing therapeutic strategies that target early disease mechanisms rather than end-stage pathology. In the present study, we investigate the impact of GSH dysregulation on neuronal-like cells under AD-relevant conditions, aiming to clarify how compromised redox buffering contributes to neurodegenerative progression.

## Materials and methods

### Reagents

All chemicals used were of analytical reagent grade. Milli-Q water was used to prepare standards and reagents. C11-Bodipy 581/591 (Cat. D3861), Hoechst33342 (Cat. H3570), were purchased from Thermo Fisher (Waltham, MA). RSL3 (Cat. B6095), Ferrostatin-1 (cat no.-A4371) and Bafilomycin A1 (Cat. A8627) were purchased from APExBIO (Boston, MA). Cycloheximide (Cat. C7698) was purchased from Sigma-Aldrich (St. Louis, MO). Liproxstatin-1 (Cat. HY-12726) was purchased from MedChemExpress (Monmouth Junction, NJ, USA). Aβ monomers (Cat No. – 641-00) were purchased from ECHELON Biosciences (Salt Lake City, UT, USA). Lactate dehydrogenase (LDH) assay kit (Cat. J2380) was purchased from Promega (Madisson, WI, USA). Iron assay kit (Cat. I7504-60) was purchased from Pointe Scientific (Canton, MI, USA). ZVAD-FMK (Cat. FMK001) was purchased from R&D Systems (Minneapolis, MN, USA). Aβ 1-42 ELISA assay kit was purchased from Thermo Fisher (Waltham, MA). Doxycycline (Cat. DSD43020-10) was purchased from dot scientific inc (Burton, MI, USA). Puromycin (Cat. BP-2956-100) was purchased from Thermo Fisher (Waltham, MA). All other chemicals were obtained from Sigma-Aldrich and Fisher Scientific. Antibodies used in this study listed in Table S1.

### Cell Culture

The human neuroblastoma cell line, SHSY-5Y cells [22, 23]was purchased from ATCC (Cat. no. CRL-2266, New York, NY, USA) and was grown in MEM:F12 media (1:1) with 15% FBS, 100 U/ml penicillin, and streptomycin at 37 in a humidified 5% CO_2_ incubator. HeLa cells were maintained in DMEM with high glucose, 10% FBS, and 100 U/ml of penicillin and streptomycin at 37 in a humidified 5% CO_2_ incubator.

### Generation of APP_695_ and APP_Swe/Ind_ Stable Overexpression Cell Lines

Wild-type APP_695_ cDNA and APP harboring the Swedish/Indiana (APP_Swe/Ind_) mutations were generously provided by Drs. Dennis Selkoe and Tracy Young-Pearse [24] (Addgene plasmids #30137 and #30145). Both APP_695_ and APP_Swe/Ind_ constructs were subcloned into the lentiviral expression vector pLVX-TetOne-Puro (Takara Bio, San Jose, CA) to enable doxycycline-inducible expression.

Lentiviral particles were produced in HEK293T cells following previously published protocols[25]. SH-SY5Y cells were infected with the respective lentiviruses for 48 hours, followed by selection with puromycin (0.5ng/mL) to generate stable overexpression cell lines. For experimental studies, cells were maintained in doxycycline-free medium. To induce APP_695_ or APP_Swe/Ind_ expression, doxycycline (300 ng/mL) was added to the culture medium, and APP expression levels were verified by immunoblot analysis.

### Generation of GCLC Knockout SH-SY5Y Cells by CRISPR/Cas9 Editing

The sgRNA targeting GCLC was delivered using the lentiCRISPR v2 vector (a gift from Dr. Feng Zhang; Addgene plasmid #52961) following our previously established protocol and validated gRNA sequences[26]. Lentiviral particles were generated in HEK293T cells by co-transfection of lentiCRISPR v2, VSV-G, BH10 φ-env-, and pRev constructs.

SH-SY5Y cells were infected with the lentiviral particles and cultured for 48 hours prior to selection. Stably transduced cells were selected with 0.5 ng/mL puromycin. For single-cell cloning, infected cells were serially diluted and plated into 96-well plates. Individual clonal cell lines with GCLC knockout were validated by immunoblot analysis.

### Preparation of A*β* oligomers

Aβ oligomers were prepared according to the method described by Stine et al [27]. Briefly, Aβ monomers were first dissolved in DMSO at a concentration of 5 mM and then diluted to 100µM in phenol red-free Ham’s F12 media. The solution was incubated at 4°C for 72 hours to allow oligomer formation before being added to the cell culture system. The diluted Aβ preparation was used immediately after the 72 -hour incubation and was not stored for future use. The formation of Aβ oligomers in each batch was validated by immunoblot analysis.

### Membrane lipid peroxidation assay using C11-Bodipy

SH-SY5Y, SH-SY5Y-APP_695,_ and SH-SY5Y-APP_swe/Ind_ cells were seeded at a density of 5×10^4^ cells per well in 24-well plates containing coverslips. Cells were treated with or without 300 ng/ml doxycycline, 5 µM Aβ, 350 µM buthionine sulfoximine (BSO), 5 µM dimethyl fumarate (DMF), and 0.1 µM ferrostatin-1 (Fer-1), as indicated.

At the end of the treatment period, C11-Bodipy 581/591 was added to a final concentration of 5 μM and incubated for 30 minutes. Coverslips were then gently washed with PBS and fixed with 4% paraformaldehyde containing 1 μg/ml Hoechst 33342 for 10 minutes at 37°C. After washing, the coverslips were mounted for confocal image acquisition.

### Malondialdehyde (MDA) assay

MDA levels were measured using the thiobarbituric acid reactive substances (TBARS) colorimetric assay kit (Cat. 10009055, Cayman Chemical, Ann Arbor, MI) following the manufacturer’s instructions. MDA values were normalized to total protein content.

### 4-hydroxynonenal (4-HNE) assay

HNE levels were assessed using multiple approaches. Immunofluorescence (IF) staining and immunoblot analysis were performed using the 4-HNE antibody (Cat. HNE11-s, Alpha Diagnostic, San Antonio, TX). SH-SY5Y, SH-SY5Y-APP_695,_ and SH-SY5Y-APP_Swe/Ind_ cells were seeded at a density of 5×10^4^ cells per well in 24-well plates containing coverslips, and treated with or without 300ng doxycycline, 5 µM Aβ, 350 µM BSO, 5 µM DMF, and 0.1 µM Fer-1, as indicated in the culture medium. After 48 hours of treatment, cells were rinsed with PBS, fixed, and processed for IF staining according to our previous report [28].

### Cell viability assay

Cells were seeded at a density of 10,000 cells per well in 96-well plates. Treatments were administered with or without 300 ng/ml Doxycycline, 5 µM Aβ, 350 µM BSO, 5 µM DMF, and 0.1 µM Fer-1, as indicated in the culture medium. At the end of the treatment period, cell viability was assessed using the CCK-8 kit (Cat. HY-K0301, MedChemExpress, Monmouth Junction, NJ) according to the manufacturer’s instructions. Cell viability was normalized relative to the cell number measured on the day immediately prior to treatment.

### LDH assay

SH-SY5Y, SH-SY5Y-APP_695,_ and SH-SY5Y-APP_Swe/Ind_ cells were seeded in 96-well plates 16 hours prior to treatment. At the end of treatment period, cultured medium was collected and diluted 1:25 in LDH storage buffer (200 mM Tris-HCl, pH 7.3, 10% glycerol, 1% BSA). LDH levels were measured using the LDH detection kit (Cat. J2380, Promega, Madison, WI, USA) following the manufacturer’s instructions. Briefly, 25 μL of samples and standards were added to a 96-well opaque plate, followed by 25 μL of LDH detection reagent. The plate was incubated for 1 hour, and luminescence was measured to determine LDH activity. LDH concentrations were calculated using a standard curve and normalized to cell number.

### Iron assay

Iron levels were measured following a previously described method [29]. In brief, SH-SY5Y, SH-SY5Y-APP_695,_ and SHSY5Y-APP_Swe/Ind_ cells were seeded at a density of 2×10^5^ cells per well in 6-well plates and treated with and without 300 ng/ml doxycycline, 5 µM Aβ, 350 µM BSO, 5 µM DMF, and 0.1 µM Fer-1, as indicated in the culture medium. At the end of the treatment period, cells were harvested, washed with PBS, and lysed by two freeze-thaw cycles (−80°C to room temperature) in 50 μL Milli-Q water. Following centrifugation, the supernatant was collected and used for the iron quantification.

### Confocal microscopy

All fluorescence images were acquired using a Leica STELLARIS confocal microscopy instrument. All images were processed using LAX-X or ImageJ software.

### Immunoblotting assay

The protein concentration was assessed using the Bicinchoninic Acid (BCA) assay (ThermoFisher), and equal amounts of protein were loaded onto SDS-PAGE gels for electrophoresis. Proteins were transferred to 0.45 μm pore size PVDF membrane (Bio-Rad) and subjected to immunoblot analysis. After primary antibody incubation, an HRP-conjugated secondary antibody was applied, and signals were detected using an enhanced chemiluminescence (ECL) substrate.

Protein loading was normalized to housekeeping control proteins, such as GAPDH, when appropriate. For treatments that affected housekeeping gene expression, total protein normalization was performed using Bio-Rad stain-free SDS-PAGE gels, and protein loading was quantified based on total lane intensity using the ChemiDoc MP imaging system (Bio-Rad).

### A*β* 1-42 ELISA

Aβ 1-42 levels were measured using a commercially available ELISA kit according to the manufacturer’s protocol. Briefly, culture medium was collected 36 hours after doxycycline-induced APP overexpression and diluted by 1:100 in the provided diluent buffer prior to being added to the ELISA plate for analysis.

### GSH Assay

GSH levels were determined using the colorimetric assay as described[30]. Briefly, cultured cells were homogenized on ice in 0.6% sulfosalicylic acid prepared in a freshly made 0.1 M potassium phosphate buffer (pH 7.5) containing 5 mM EDTA. The homogenates were centrifuged, and the resulting supernatants were used for the GSH assay. GSH levels were quantified using a glutathione reductase (GR)-dependent enzymatic recycling assay with β-NADPH following derivatization with 5,5′-dithiobis-(2-nitrobenzoic acid) (DTNB).

### Statistical analysis

All data are presented as mean ± standard deviation (SD), and all statistical analyses were conducted using GraphPad Prism 10.6.0 software. Comparisons between two groups were performed using a two-tailed Student’s *t*-test, while multi-group comparisons were analyzed by one-way ANOVA, as described in the figure legends. A *P*-value <0.05 was considered statistically significant. Statistical significance is indicated as follows: \**P* < 0.05, \*\**P* < 0.01, \*\*\**P* < 0.001, \*\*\*\**P* < 0.0001.

## Result

### GSH depletion exacerbates cell death in APP Swedish/Indiana mutant-overexpressing cells

To investigate the impact of GSH dysregulation on Aβ-associated pathogenesis, we generated doxycycline-inducible SH-SY5Y cell lines overexpressing either wild-type APP (APP_695_) or APP carrying the Swedish/Indiana mutations (APP_Swe/Ind_) using a lentiviral system. Treatment with 300 ng/ml doxycycline robustly induced APP_695_ and APP_Swe/Ind_ expression, as confirmed by immunoblot analysis (Figure 1A). As expected, APP_Swe/Ind_ overexpression resulted in a marked increase in Aβ 1-42 secretion after 48 hours of inducing overexpression, approximately 900-fold higher than APP695, as measured by ELISA (Figure 1B).

**Figure 1.**
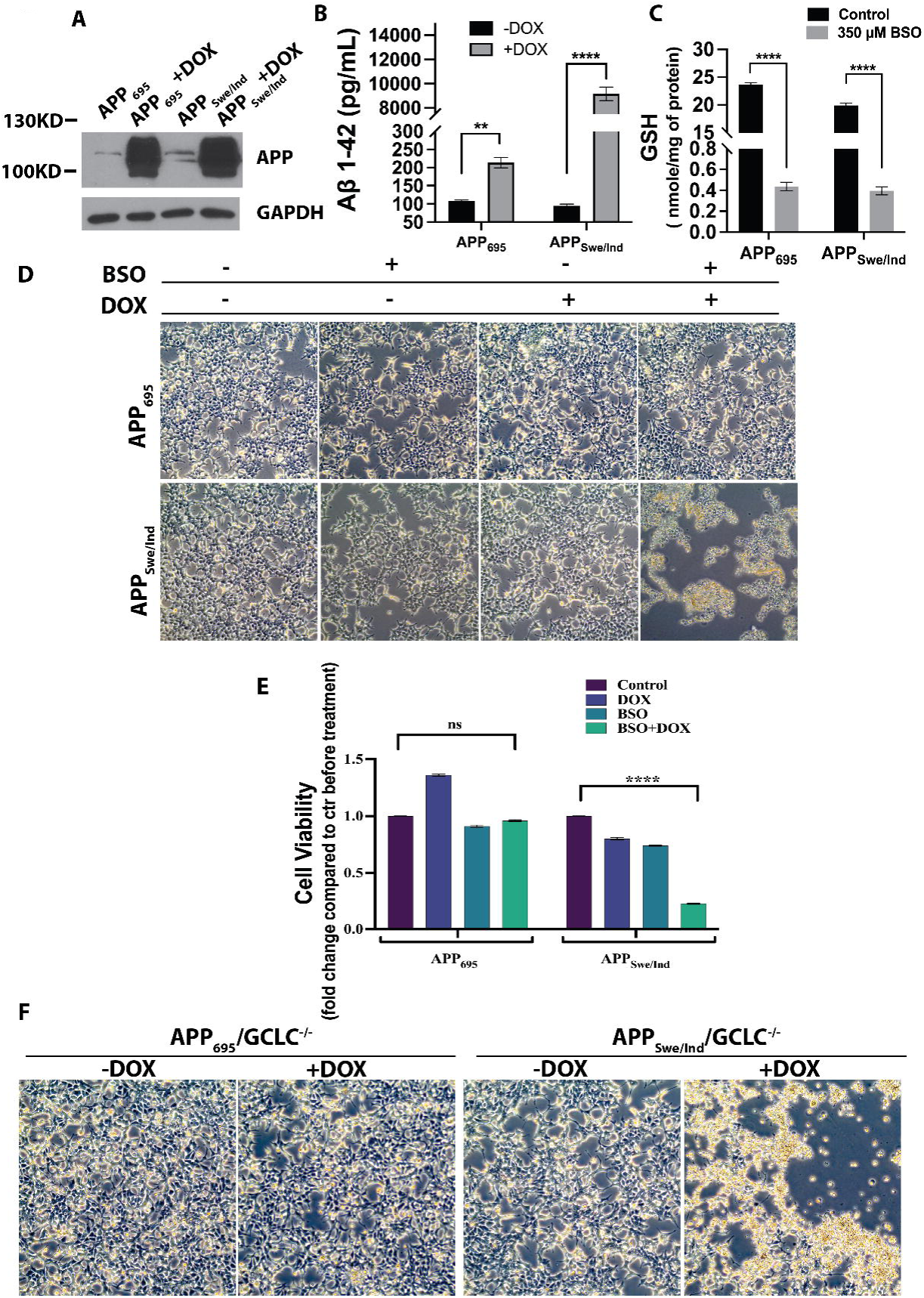
Glutathione dysregulation in APP_Swe/Ind_ overexpressing cells causes cell death. A) Immunoblot showed doxycycline induced overexpression of APP_695_ and APP_Swe/Ind_ in SH-SY5Y cells. B) ELISA for Aβ1-42 shows SH-SY5Y cells overexpressing APP_Swe/Ind_ is generating high level of Aβ1-42 in the cell culture system. C) 350µM of BSO depleted GSH in both APP_695_ and APP_Swe/Ind_ overexpressing cells. D) Brightfield images showing doxycycline induced APP_Swe/Ind_ overexpressing cells when treated with BSO has morphological changes after 36 hours of treatment. E) CCK8 analysis showing the cell viability of APP_Swe/Ind_ overexpressing cells after 36 hours of BSO treatment decreases compared to APP_695_ overexpressing cells. F) Brightfield images showing cell death of APP_Swe/Ind_ overexpressing SH-SY5Y cells after GCLC knockdown. Ordinary one-way ANOVA with Tukey’s multiple comparisons test and an unpaired t-test was conducted accordingly. Significance was considered at P < 0.05, denoted as *<0.05, **<0.01, ***<0.001, ****<0.0001.

To deplete intracellular GSH, cells were treated with BSO, an inhibitor of glutamyl-cysteine ligase (GCL), the rate-limiting enzyme in de novo GSH synthesis. Dose-response testing in APP_695_ and APP_Swe/Ind_ stable cell lines without doxycycline induction demonstrated that 350 µM BSO significantly reduced intracellular GSH levels for up to 72 hours without causing detectable cytotoxicity (Figure 1C). Therefore, 350 µM BSO was used in subsequent experiments.

Upon doxycycline-induced overexpression of APP_695_ or APP_Swe/Ind_ in the presence or absence of BSO, pronounced cell death was observed in APP_Swe/Ind_ -overexpressing cells treated with 350 µM BSO for 36 hours. In contrast, APP_695_-overexpressing cells showed no significant cell death under the same conditions (Figure 1D). These findings were further confirmed by the CCK-8 viability assay. As shown in Figure 1E, BSO treatment resulted in more than 25% cell loss in APP_Swe/Ind_ -overexpressing cells compared to APP_695_-overexpressing cells at 36 hours.

For further validation, GCLC, the catalytic subunit of GCL, was knocked out by the CRISPR/Cas9 system in both the APP_695_ and APP_Swe/Ind_-overexpressing cells, and the knocked-out cells were supplemented with N-acetyl Cysteine (NAC). When APP_Swe/Ind_ was overexpressed in the GCLC^-/-^ cells, the cells started to show morphological changes 72 hours onwards, and after 100 hours marked cell death was observed. Whereas APP_695_/GCLC^-/-^ cells did not show any morphological changes under the microscope or cell death phenomenon after 100 hours.

### Intracellular GSH depletion and the APP Swedish/Indiana mutation synergistically induce lipid peroxidation

To elucidate the mechanism of cell death, we first assessed lactate dehydrogenase (LDH) release as an indicator of plasma membrane damage. As shown in Figure 2A, LDH levels in the culture medium were markedly increased in APP_Swe/Ind_-overexpressing cells treated with BSO compared to APP_695_ cells under the same conditions. Notably, LDH release was significantly attenuated in the presence of 1 µM ferrostatin-1 (Fer-1), a lipid peroxidation inhibitor, suggesting that plasma membrane damage was likely mediated by lipid peroxidation.

**Figure 2.** Lipid peroxidation in glutathione dysregulated SH-SY5Y cells overexpressing APP_695_ and APP_Swe/Ind._ A) LDH level increased in APP_Swe/Ind_ overexpressing cells treated with BSO for 36 hours compared to APP_695_ overexpressing cells treated with BSO. B) C11-BODIPY staining of BSO-treated APP_Swe/Ind_ overexpressing cells showed an increased level of membrane lipid peroxidation compared to APP_695_ overexpressing cells. C) Quantification of C-11 BODIPY staining. D) Increased level of MDA in BSO treated APP_Swe/Ind_ overexpressing cells compared to control groups and APP_695_ overexpressing cells. E) 4-HNE immunoblot shows increased level of 4-HNE modified proteins in BSO treated APP_Swe/Ind_ overexpressing cells. F) 4-HNE IF of BSO treated APP_Swe/Ind_ overexpressing cells shows an increase in 4-HNE fluorescence intensity. G) Quantification of 4-HNE IF of APP_Swe/Ind_ overexpressing cells following BSO treatment. Ordinary one-way ANOVA with Tukey’s multiple comparisons test was conducted. Significance was considered at P < 0.05, denoted as *<0.05, **<0.01, ***<0.001, ****<0.0001.

To further evaluate membrane lipid peroxidation, C11-BODIPY 581/591 staining was performed. In this assay, lipid peroxidation shifts fluorescence emission from red (∼590 nm, reduced form) to green (∼510 nm, oxidized form). As shown in Figures 2B and 2C, APP_Swe/Ind_-overexpressing cells treated with BSO exhibited a significant increase in plasma membrane lipid peroxidation compared to APP_695_ cells. Co-treatment with Fer-1 markedly reduced the oxidized lipid signal. In contrast, no significant lipid peroxidation was observed in APP_695_-overexpressing cells treated with BSO alone.

Lipid peroxidation was further validated by measuring malondialdehyde (MDA). As shown in Figure 2D, MDA level was significantly elevated in APP_Swe/Ind_-overexpressing cells treated with BSO compared to APP_695_-overexpressing cells. Moreover, both 4-HNE immunoblot and immunostaining showed a significant increase in 4-HNE level in the APP_Swe/Ind_-overexpressing cells treated with BSO, further validating increased lipid peroxidation (Figure 2E-2G, Supplementary Figure 1).

### Intracellular GSH depletion and APP Swedish/Indiana mutation synergistically induce ferroptosis in SH-SY5Y cells

To determine the mechanism underlying cell death, APP_Swe/Ind_ -overexpressing cells were co-treated with BSO and either pan-caspase inhibitor Z-VAD-FMK or the ferroptosis inhibitors ferrostatin-1 (Fer-1) and liproxstatin-1 (Lip-1), to assess whether cell death occurred through apoptosis or ferroptosis.

As shown in Figures 3A and 3B, Z-VAD-FMK failed to rescue BSO-plus APP_Swe/Ind_-induced cell death, as assessed by both cell morphology and CCK8 viability assay. Immunoblot analysis of apoptosis markers further demonstrated no change in pro-caspase 3 levels and no detectable cleaved caspase 3 (Figure 3C), confirming that apoptosis was not the primary mechanism of cell death. We used SH-SY5Y cells treated with 2.5 µM staurosporine for 4 hours as a positive control for SH-SY5Y apoptosis. In contrast, both Fer-1 and Lip-1 effectively blocked cell death induced by BSO and APP_Swe/Ind_ overexpression, as evidenced by improved cell morphology (Figure 3A) and restored viability in the CCK8 assay (Figure 3B). These findings indicate that intracellular GSH depletion in APP_Swe/Ind_-overexpressing SH-SY5Y cells triggers ferroptotic cell death.

**Figure 3.**
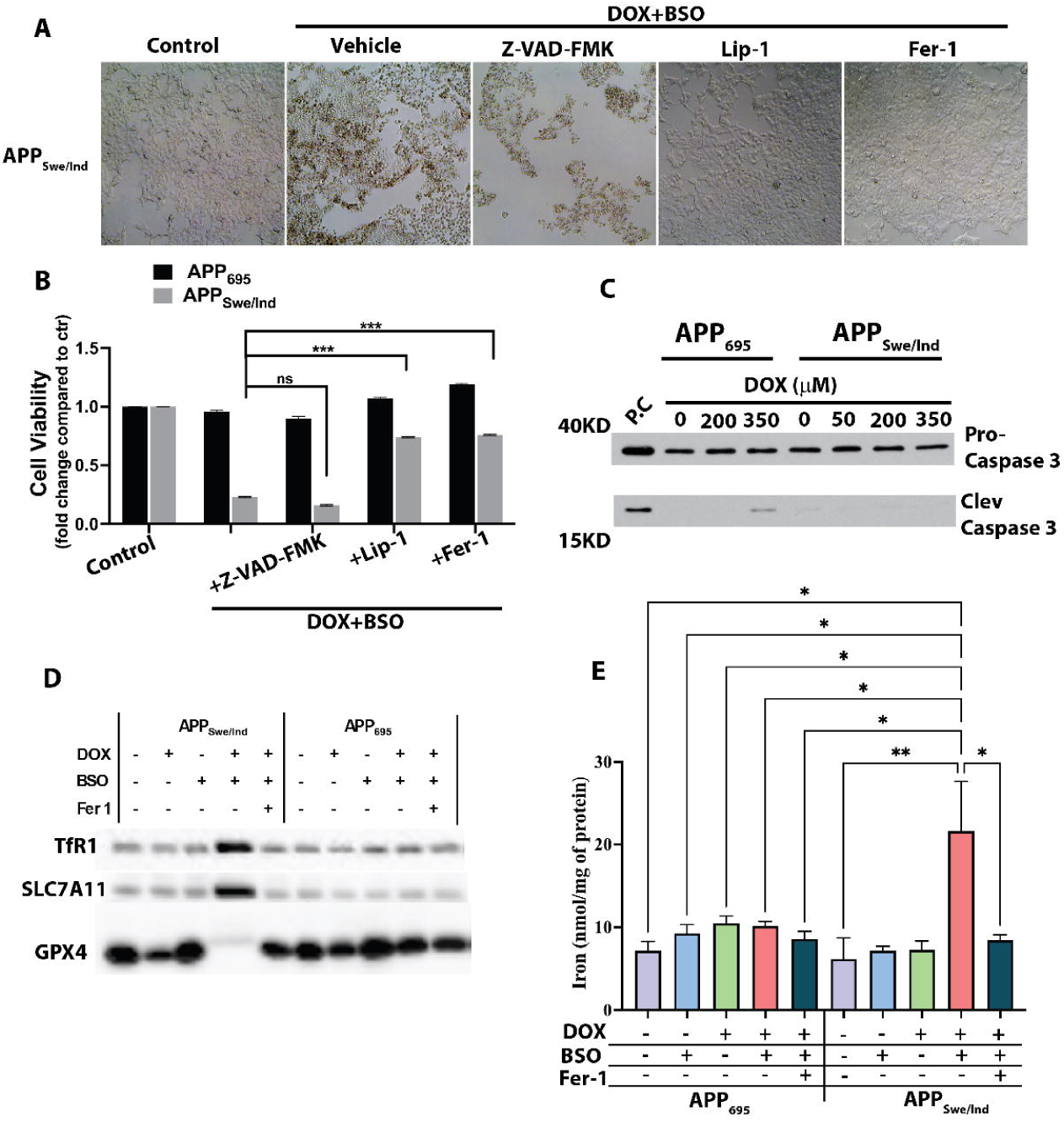
APP_Swe/Ind_-overexpressing cells treated with BSO cause ferroptosis-mediated cell death. A) Brightfield images showing cell morphology of doxycycline-induced APP_Swe/Ind_ overexpressing cells change after 36 hours of BSO treatment and rescued by Lip-and Fer-1 but not by Z-VAD-FMK. B) CCK8 assay of doxycycline induced SH-SY5Y-APP_Swe/Ind_ cells treated with BSO for 36 hours showed a decrease in cell viability and Z-VAD-FMK could not rescue the cells but Lip-1 and Fer-1 rescued the cells. No changes in cell viability were observed in SH-SY5Y-APP_695_ overexpressing cells with the same treatment. C) Immunoblot shows no change in procaspase 3 or any increase in cleaved caspase 3 after 36 hours of BSO treatment on doxycycline-induced SH-SY5Y-APP_Swe/Ind_ or -APP_695_ cells. D) Immunoblot of BSO treated and doxycycline induced SH-SY5Y-APP_Swe/Ind_ and -APP_695_ cells shows increase in TfR1 and SLC7A11 and a decrease in GPX4. E) Level of intracellular iron in BSO treated and doxycycline induced SH-SY5Y-APP_Swe/Ind_ cells increased compared to control groups and SH-SY5Y-APP_695_ cells. Ordinary one-way ANOVA with Tukey’s multiple comparisons test was conducted. Significance was considered at P < 0.05, denoted as *<0.05, **<0.01, ***<0.001, ****<0.0001.

We next examined canonical ferroptosis-related markers. Immunoblot analysis revealed a significant increase in transferrin receptor 1 (TfR1) levels (Figures 3D, Supplementary Figure 2A and B) in APP_Swe/Ind_-overexpressing cells treated with BSO. This increase was markedly attenuated by ferrostatin-1 (Fer-1) treatment. In contrast, APP_695_-overexpressing SH-SY5Y cells treated with BSO did not exhibit any significant change in TfR1 expression. The expression was compared based on the total protein load in the stain-free gel (Supplementary Figure 2A). TfR1 expression in SH-SY5Y-APP_Swe/Ind_ cells was further validated by immunofluorescence staining (Supplementary Figures 2C and D).

The cystine/glutamate antiporter SLC7A11, which is essential for intracellular cysteine uptake and GSH synthesis, showed minimal change in SH-SY5Y cells treated with 350 µM BSO alone. However, in APP_Swe/Ind_-overexpressing cells treated with BSO, SLC7A11 expression increased more than ninefold (Figures 3D, and Supplementary Figures 2A and E). No comparable change was observed in SH-SY5Y-APP_695_ cells following BSO treatment (Figure 3D and Supplementary Figure 2E).

GPX4, a key enzyme that protects against lipid peroxidation and a central regulator of ferroptosis, was significantly reduced in SH-SY5Y-APP_Swe/Ind_ cells and further decreased upon BSO treatment (Figure 3D and Supplementary Figure 2A and F). In contrast, SH-SY5Y-APP_695_ cells showed no significant change under the same conditions. Interestingly, BSO treatment alone increased GPX4 levels in both cell types, likely reflecting a compensatory response to oxidative stress.

Because transferrin receptor 1 (TfR1) mediates iron uptake, we also assessed intracellular iron levels. BSO treatment did not significantly alter iron content in APP_695_-overexpressing cells compared to controls (Figure 3E). However, APP_Swe/Ind_-overexpressing cells treated with BSO exhibited a significant increase in intracellular iron levels relative to controls (Figure 3E).

Together, these data demonstrate that intracellular GSH depletion synergizes with the APP_Swe/Ind_ mutation to promote ferroptosis in SH-SY5Y cells.

### A combination of Aβ oligomers and intracellular GSH depletion synergistically induces lipid peroxidation and ferroptosis in SH-SY5Y cells

Because APP_Swe/Ind_-overexpressing cells secrete substantial amounts of Aβ, we reasoned that the observed cell death may result from the combined effects of accumulated Aβ and intracellular GSH dysregulation. Aβ oligomers are well-recognized cytotoxic species capable of triggering neurodegeneration; therefore, we hypothesized that intracellular GSH depletion would sensitize cells to Aβ oligomer-mediated toxicity.

The Aβ oligomers prepared for this study were predominantly octamers, as confirmed by immunoblot analysis (Supplementary Figure 3A). We first evaluated the tolerance of SH-SY5Y cells to Aβ oligomers alone and observed no noticeable cell death or morphological changes at or below 5 µM Aβ treatment for 48 hours (Supplementary Figure 3B). To deplete intracellular GSH, we used dimethyl fumarate (DMF), which rapidly reduces GSH levels through direct conjugation[31]. As shown in Figure 4A, 5 µM DMF significantly decreased intracellular GSH in SH-SY5Y cells without inducing detectable cell death for up to 24 hours. Based on these findings, 5 µM Aβ and 5 µM DMF were used for subsequent experiments.

**Figure 4.**
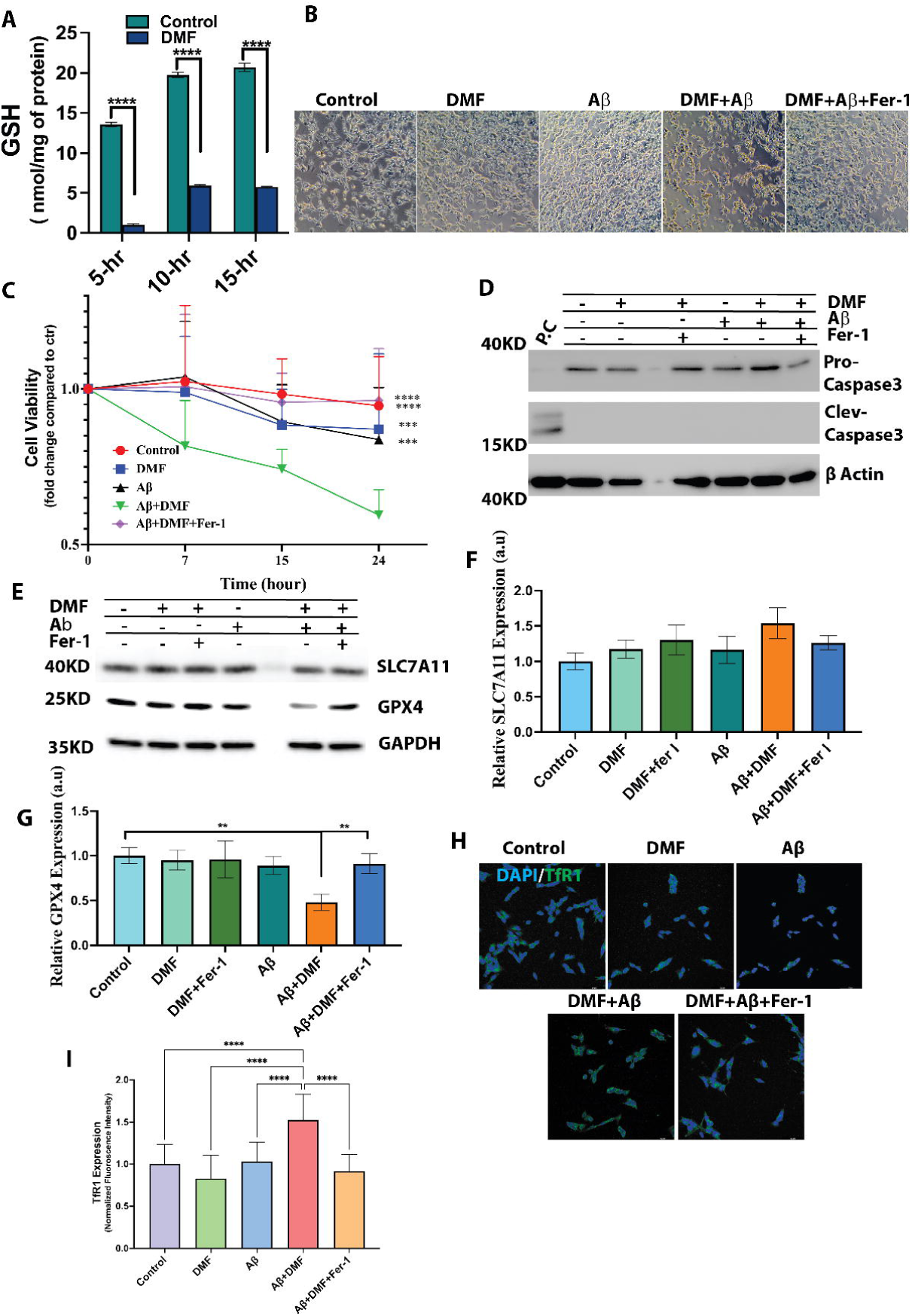
Synergistic treatment of Aβ and DMF effects canonical markers of ferroptosis. A) GSH was depleted significantly in SH-SY5Y cells by 5µM DMF till 15 hours. B) Brightfield images showing the cellular morphology of SH-SY5Y cells change when treated with Aβ and DMF for 7 hours. C) CCK8 cell viability of SH-SY5Y cells shows a decrease in cell viability following Aβ and DMF treatment from 7 hours onward. D) Immunoblot shows no change in procaspase 3 or any increase in cleaved caspase 3 after Aβ and DMF cotreatment of SH-SY5Y cells. E-G) SH-SY5Y cells had an increase in SLC7A11 and a decrease in GPX4 when treated with Aβ and DMF compared to when SH-SY5Y cells were treated with either Aβ or DMF. Fer-1 brought back the level of both SLC7A11 and GPX4 to the base level. H) TfR1 IF shows an increase in fluorescence intensity in SH-SY5Y cells treated with Aβ and DMF for 15 hours and decreased back to base level by Fer-1. I) Quantification of of TfR1 immunofluorescence intensity. Ordinary one-way ANOVA with Tukey’s multiple comparisons test and an unpaired t-test was conducted accordingly. Significance was considered at P < 0.05, denoted as *<0.05, **<0.01, ***<0.001, ****<0.0001.

As shown in Figure 4B, treatment with 5 µM DMF or 5µM Aβ alone did not induce significant cell death for up to 48 hours. In contrast, co-treatment with 5 µM DMF and 5µM Aβ oligomers resulted in marked cell death as early as 7 hours (Figure 4B). This effect was significantly attenuated by co-treatment with Fer-1 (Figure 4B). CCK8 assays further confirmed these observations: cell viability progressively declined beginning at 7 hours and decreased to approximately 60% at 24 hours following combined DMF and Aβ treatment. Importantly, Fer-1 restored viability to nearly 100% (Figure 4C).

Immunoblot analysis showed no changes in pro-caspase-3 level and no detectable cleaved caspase-3 after 15 hours of combined treatment (Figure 4D), suggesting that apoptosis was not involved. Immunoblot analysis showed GPX4 levels were significantly decreased, and SLC7A11 levels were modestly increased in Aβ and DMF co-treated cells compared with controls (Figure 4E-4G). Immunofluorescence staining further demonstrated upregulation of TfR1 in SH-SY5Y cells following the combination of Aβ and DMF treatment (Figure 4H, 4I, Supplementary figure 4).

Oxidized lipids in the plasma membrane were assessed using BODIPY 581/591 C-11. Co-treatment with DMF and Aβ oligomers resulted in significantly increased oxidized lipid levels and a corresponding decrease in reduced lipid levels in the plasma membrane compared with control groups (Figure 5A and Supplementary Figure 5). Importantly, Fer-1 again markedly reduced oxidized lipid levels while restoring reduced lipid levels (Figure 5A and Supplementary Figure 5).

**Figure 5.**
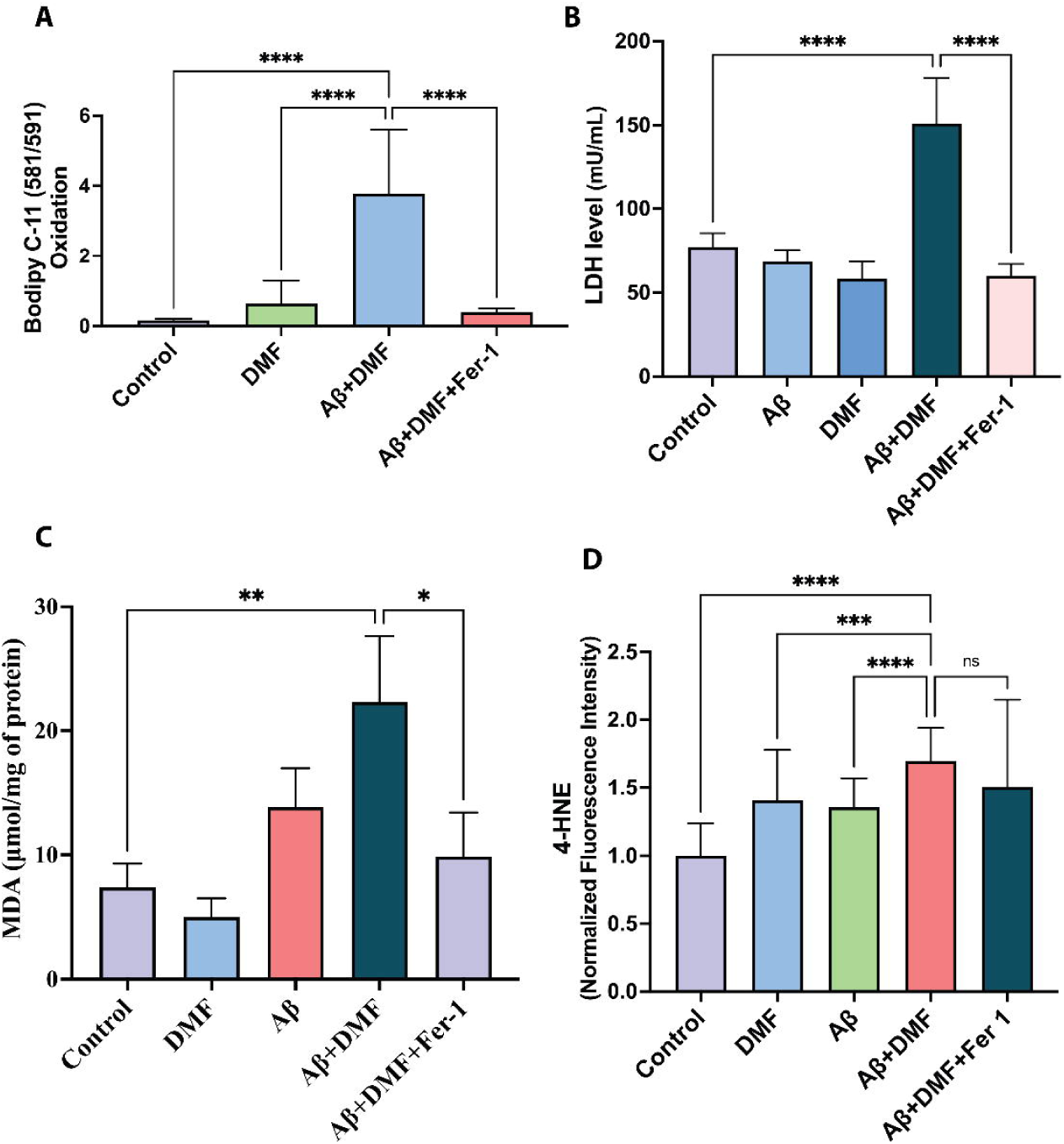
GSH depletion induces lipid peroxidation in presence of Aβ peptide. A) Quantification of C11-BODIPY 581/591 staining in SH-SY5Y cells treated with Aβ and DMF for 15 hours showed increased membrane lipid peroxidation which was rescued by Fer-1. B) LDH level significantly increased in SH-SY5Y cells treated with Aβ and DMF for 15 hours. C) MDA level significantly increased in SH-SY5Y cells treated with Aβ and DMF for 15 hours. D) Quantification of 4-HNE immunofluorescence staining of SH-SY5Y cells co-treated with Aβ and DMF for 15 hours showed increased formation of 4-HNE. Ordinary one-way ANOVA with Tukey’s multiple comparisons test was performed. Significance was considered at P < 0.05, denoted as *<0.05, **<0.01, ***<0.001, ****<0.0001.

To further evaluate membrane lipid damage, we checked LDH release. Co-treatment with DMF and Aβ oligomers caused the cells to release significantly higher levels of LDH compared to the controls, and this effect was reversed by Fer-1 (Figure 5B). Additionally, analysis of other lipid peroxidation markers, MDA and 4-HNE, showed elevated levels in cells treated with Aβ and DMF compared to control groups (Figure 5C-5D, Supplementary figure 6), and were alleviated by Fer-1, which further confirms increased lipid oxidation.

Finally, we also checked the iron content of Aβ-and DMF co-treated SH-SY5Y cells and found that, under the treatment conditions, the iron content increased in the SH-SY5Y cells to about 20nM/mg protein, compared with ∼12 nM/mg in control cells (Figure 5H).

### Combination of Aβ oligomers and intracellular GSH depletion activates autophagy

Several studies have suggested that autophagy-mediated GPX4 degradation can contribute to ferroptosis[32]. In our study, we observed significant suppression of GPX4 expression in SH-SY5Y cells under conditions of intracellular GSH depletion combined with elevated Aβ levels, either through APP_Swe/Ind_ overexpression or exogenous Aβ oligomer treatment. Therefore, we examined the status of autophagy under these conditions. As shown in Figures 6A and 6B (and Supplementary Figure 2A), immunoblot analysis confirmed a significant elevation of LC3-II in APP_Swe/Ind_-overexpressing SH-SY5Y cells following BSO treatment. A similar increase in LC3-II was observed in SH-SY5Y cells treated with DMF and Aβ in combination (Figure 6C, 6D).

**Figure 6.**
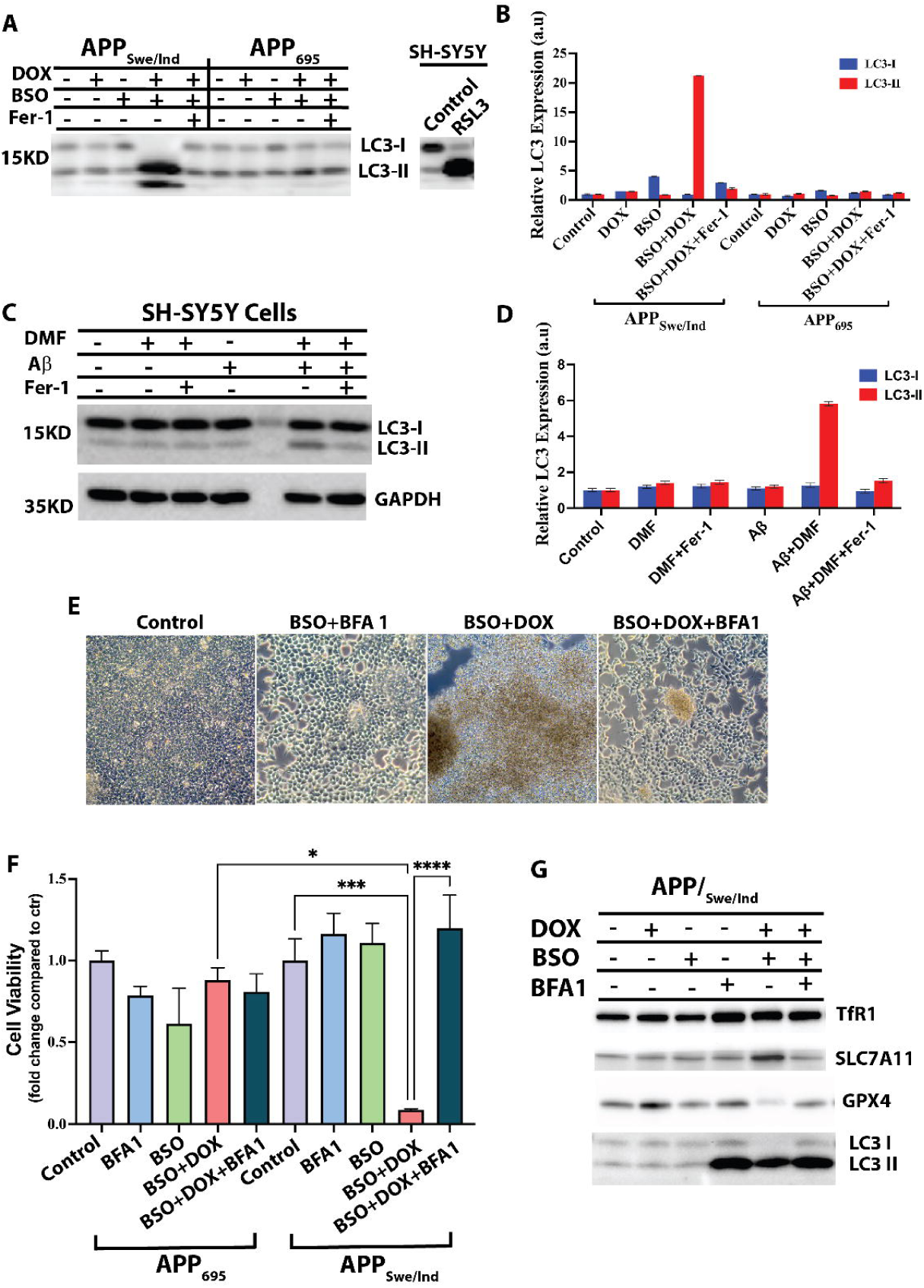
Aβ and GSH depletion instigates autophagy driven ferroptosis. A-B) Immunoblot of LC3 in APP_Swe/Ind_ and APP_695_ overexpressing cells treated with BSO for 36 hours. 2.5 µM RSL3-treated SH-SY5Y cells were used as a comparable control for ferroptosis. LC3-II levels increased in APP_Swe/Ind_-overexpressing cells treated with BSO. C-D) Immunoblot of LC3 in SH-SY5Y cells treated with Aβ and DMF for 15 hours, showing an increase in LC3-II level. E) Brightfield images showing cellular morphology of APP_Swe/Ind_ overexpressing cells treated with BSO and BFA 1 remain the same as the control after 48 hours of treatment. F) CCK8 cell viability of APP_Swe/Ind_ and APP_695_ overexpressing cells treated with BSO and BFA 1 after 48 hours of treatment showing APP_Swe/Ind_ overexpressing cells treated with BSO were rescued by BFA 1 . G-K) Immunoblot of TfR1, SLC7A11, GPX4, and LC3 in APP_Swe/Ind_ overexpressing cells treated with BSO and BFA 1 for 36 hours had the level rescued back to the base level. Ordinary one-way ANOVA with Tukey’s multiple comparisons test and an unpaired t-test was conducted accordingly. Significance was considered at P < 0.05, denoted as *<0.05, **<0.01, ***<0.001, ****<0.0001.

To further investigate the role of autophagy in Aβ- and GSH depletion-induced ferroptosis, we utilized bafilomycin A1 (BFA1), a potent inhibitor of autophagy that blocks autophagosome-lysosome formation[33]. We hypothesized that inhibition of autophagy would restore GPX4 expression, thereby blocking ferroptosis and rescuing cells undergoing ferroptotic cell death. Accordingly, BFA1 was added 24 hours after initiating BSO treatment in APP_Swe/Ind_-overexpressing SH-SY5Y cells. APP_695_ -overexpressing cells were treated in parallel as controls.

Notably, APP_Swe/_Ind-overexpressing cells were protected from BSO-induced cell death in the presence of BFA1 (Figure 6E, Supplementary figure 7A). CCK8 further confirmed this observation, showing a dramatic increase in cell viability from approximately 10% under BSO treatment to close to 100% with BFA1 co-treatment (Figure 6F). As expected, immunoblot analysis showed that GPX4 protein expression was restored and TfR1 and SLC7A11 protein expression was back to baseline in cells co-treated with BSO and BFA1 compared with BSO alone (Figure 6G, Supplementary Figure 7B).

Collectively, these results support the conclusion that autophagy contributes to Aβ-associated ferroptosis under conditions of GSH dysregulation and that inhibition of autophagy can rescue cells from ferroptotic cell death.

## Discussion

Aging is the primary risk factor for AD [34]. Several hallmarks of brain aging, including increased oxidative stress and elevated Aβ accumulation, are closely associated with neurodegenerative processes [35]. It is widely accepted that the aged brain environment either initiates or promotes the molecular events underlying AD [34]. In the present study, we sought to model neuronal-like cell death under conditions that mimic aspects of brain aging. To achieve this, we established a cellular environment characterized by increased Aβ production/presence and reduced intracellular GSH levels. Our findings demonstrate that the combination of Aβ accumulation and GSH depletion synergistically induces lipid peroxidation, plasma membrane damage, and ferroptosis-mediated cell death. Mechanistically, the enhanced lipid peroxidation is likely driven by suppression of GPX4 protein expression through an autophagy-mediated degradation pathway when cells are exposed to both Aβ and GSH depletion.

Multiple studies have found an aging-associated decrease in systemic [36–39] and brain GSH levels [17, 18, 40, 41], which is further exacerbated in patients with mild cognitive impairment and AD [3, 19, 20, 42]. Moreover, it was found that GSH depletion in the brain causes major neurodegeneration and cognitive deficits in animal models [43, 44]. GSH is the endogenous antioxidant, which is required by GPX4 to mitigate the effects of oxidative stress by reducing the oxidized polyunsaturated fatty acids (PUFA) [45, 46]. Because of this, depletion of GSH is considered the primary step of ferroptosis [47]. On the other hand, GSH dysregulation is also known for its effect on APP metabolism by affecting BACE1 activity [14, 48].

Ferroptosis is an iron-dependent regulated cell death mechanism, which can be induced by lipid peroxidation of membrane PUFAs [49]. Increased lipid peroxidation is a prominent feature of brain aging and contributes to heightened vulnerability to neurodegeneration. Jove et al. [50] emphasized that aging brains exhibit region-specific lipid alterations and accumulation of lipoxidative damage, with mitochondrial ATP synthase identified as a key target, linking oxidative lipid injury to impaired energy metabolism. Expanding this concept in the context of AD, Majernikova et al.[51] demonstrated reduced expression of GPX4 and activation of ferroptosis pathways in postmortem human brain tissue, accompanied by enhanced lipid peroxidation. Together, these findings support the view that dysregulated lipid redox homeostasis is one of the central drivers of brain aging and AD progression [47].

Accumulating evidence indicates that catalytic transition metals play a central role in regulating ferroptotic-mediated cell death. Under physiological conditions in young, healthy individuals, iron and copper are tightly regulated and securely bound by specialized proteins, including transferrin and ferritin for iron, as well as various cuproenzymes for copper[52, 53]. This stringent sequestration limits the availability of redox-active, labile metal ions, thereby minimizing metal-catalyzed lipid peroxidation.

With aging, however, metal homeostasis becomes progressively dysregulated. Both iron and copper accumulate in tissues such as the brain[54], increasing the pool of redox-active metal ions. Elevated levels of these metals have been repeatedly documented in neurodegenerative disorders. For example, increased redox-active iron and copper are enriched in amyloid plaques and neurofibrillary tangles in AD [55]. Such metal accumulation creates a pro-oxidative environment that favors lipid peroxidation-mediated neurodegeneration through ferroptosis.

In the present study, we modeled aging brain-like conditions in SH-SY5Y cells by increasing Aβ levels and reducing intracellular GSH. Under these conditions, we observed marked lipid peroxidation, indicating that Aβ oligomers promote neuronal plasma membrane damage when GSH is depleted, a scenario consistent with reports from prodromal and AD patients[56]. To reduce intracellular GSH, we employed both pharmacological and genetic approaches. Specifically, we used BSO to inhibit GCL activity [57], DMF to directly conjugate and deplete GSH [58], and genetic deletion of GCLC to block de novo GSH synthesis. In all cases, the combination of GSH depletion and Aβ oligomers synergistically induced SH-SY5Y cell death through ferroptosis.

We observed significant alterations in ferroptosis-related gene expression following combined treatment with Aβ oligomers and GSH depletion. Notably, TfR1, a key iron-transport protein responsible for cellular iron uptake [59], was markedly upregulated. Aberrant TfR1 elevation likely increases intracellular redox-active iron, thereby promoting lipid peroxidation and ferroptotic cell death. Consistent with the known association between iron accumulation and Alzheimer’s disease (AD) [60], we also detected increased intracellular iron levels under both experimental approaches, providing a mechanistic rationale for TfR1 upregulation during ferroptosis in the presence of Aβ and GSH deficiency.

We further found that the glutamate-cystine antiporter SLC7A11 was significantly upregulated during chronic GSH dysregulation with Aβ exposure. SLC7A11 plays a critical role in maintaining intracellular GSH levels by importing cystine for GSH synthesis [61]; thus, its upregulation likely represents a compensatory response to elevated oxidative stress. It was found in the postmortem AD brain that there was an increase in the light chain of SLC7A11 [62], and a similar increase was observed in an AD mouse model [63]which is consistent with our current findings. Collectively, these findings align with previous observations of increased iron deposition and altered SLC7A11 expression in AD[61] and strengthen the link between Aβ toxicity, GSH dysregulation, and ferroptotic signaling.

Importantly, we observed a significant downregulation of GPX4 protein following combined treatment with Aβ oligomers and GSH depletion. GPX4 is the only enzyme capable of directly detoxifying lipid peroxides, and its activity requires GSH as a cofactor[61, 64]. Impairment of GPX4 function is a central event in the canonical ferroptosis pathway. Consistent with our findings, reduced GPX4 levels have been reported in post-mortem AD brains [51, 65], and genetic ablation of Gpx4 in mice leads to cognitive decline and neurodegeneration [66]. We therefore speculate that in the aging brain, where GSH dysregulation and Aβ accumulation coexist, suppression of GPX4 may promote neuronal degeneration through lipid peroxidation and ferroptotic cell death.

Interestingly, our data suggest that decreased GPX4 protein levels are mediated, at least in part, by autophagy-dependent degradation. This is supported by our observation that co-treatment with bafilomycin A1 (BFA1), which blocks autophagosome-lysosome fusion, almost completely rescued SH-SY5Y cells from ferroptosis. Our findings are in good agreement with previous studies demonstrating that autophagy contributes to ferroptosis initiation. For example, Wu et al. reported that chaperone-mediated autophagy participates in ferroptosis induction [67], and Lee et al. showed that autophagy enhances ROS-mediated ferroptotic cell death [68].

In this study, we comprehensively examined the interplay between GSH dysregulation and Aβ and demonstrated that GSH depletion in the presence of the Aβ peptide triggers ferroptotic cell death. Our findings underscore the critical importance of maintaining redox homeostasis and adequate GSH levels in the AD brain. Furthermore, our data suggest that by autophagic flux may represent a potential therapeutic strategy to prevent ferroptosis-mediated neurodegeneration in AD.

## Author contributions

XF conceived the research; KR, CH, and ZW acquired the data; XF supervised the research; KR, CH, ZW, and XF analyzed and interpreted the data; KR and XF wrote the manuscript. All authors critically reviewed and provided input to the manuscript.

## Supporting information

Antibodies and their resource, dilution, and application

## Acknowledgments

This research was supported by grants from EY028158(XF), EY032488 (XF), the James Fickel Alzheimer’s Disease Research Fund MCGFD01043 (XF), and NEI Center Core Grant for Vision Research (P30EY031631) at Augusta University.

