## Supplementary figures and images for "β-Amyloid and Glutathione Dysregulation Cooperatively Drive Lipid Peroxidation and Ferroptosis in Neuron-Like Cells"

### Antibodies and their resource, dilution, and application

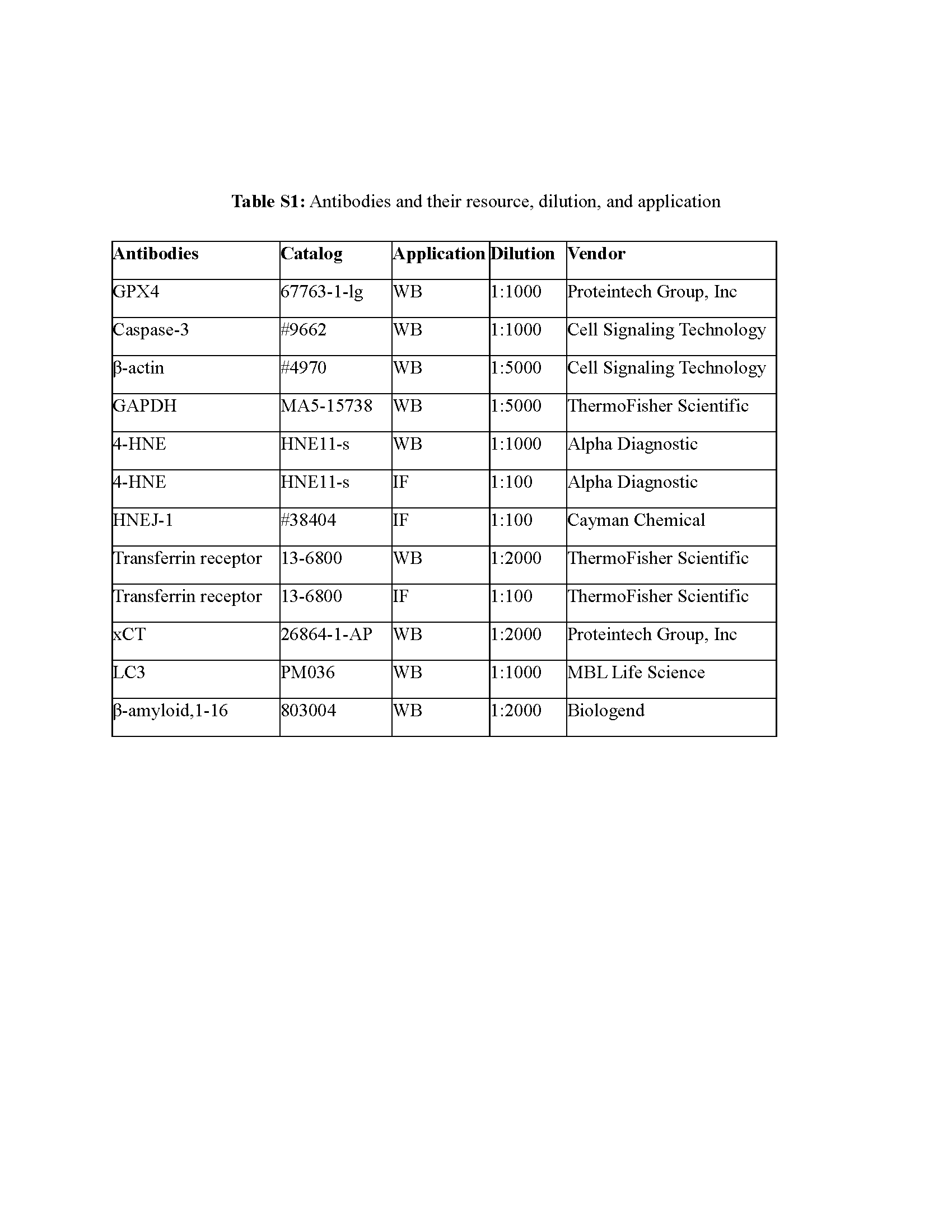

### Supplementary figure 1

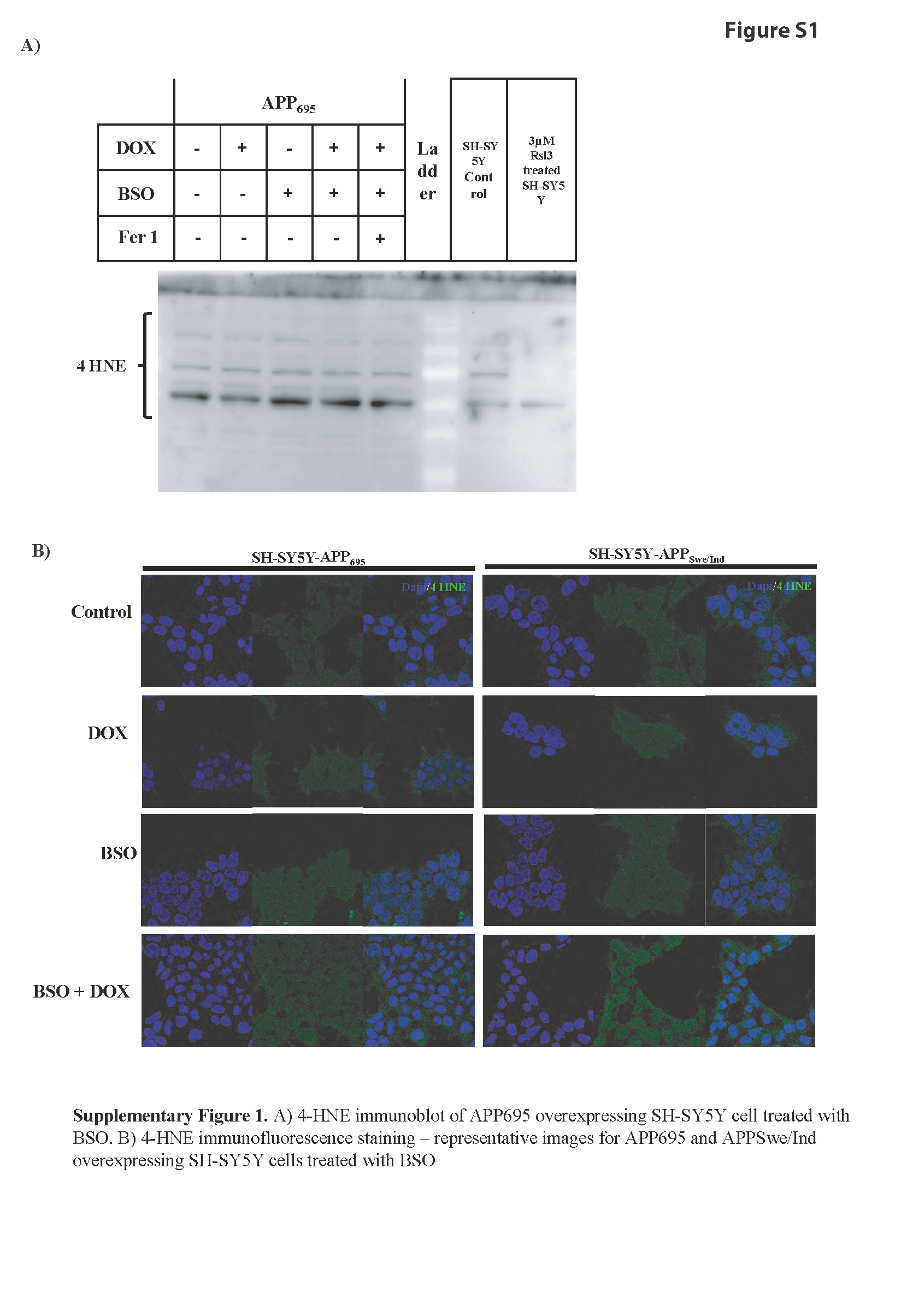

### Supplementary figure 2

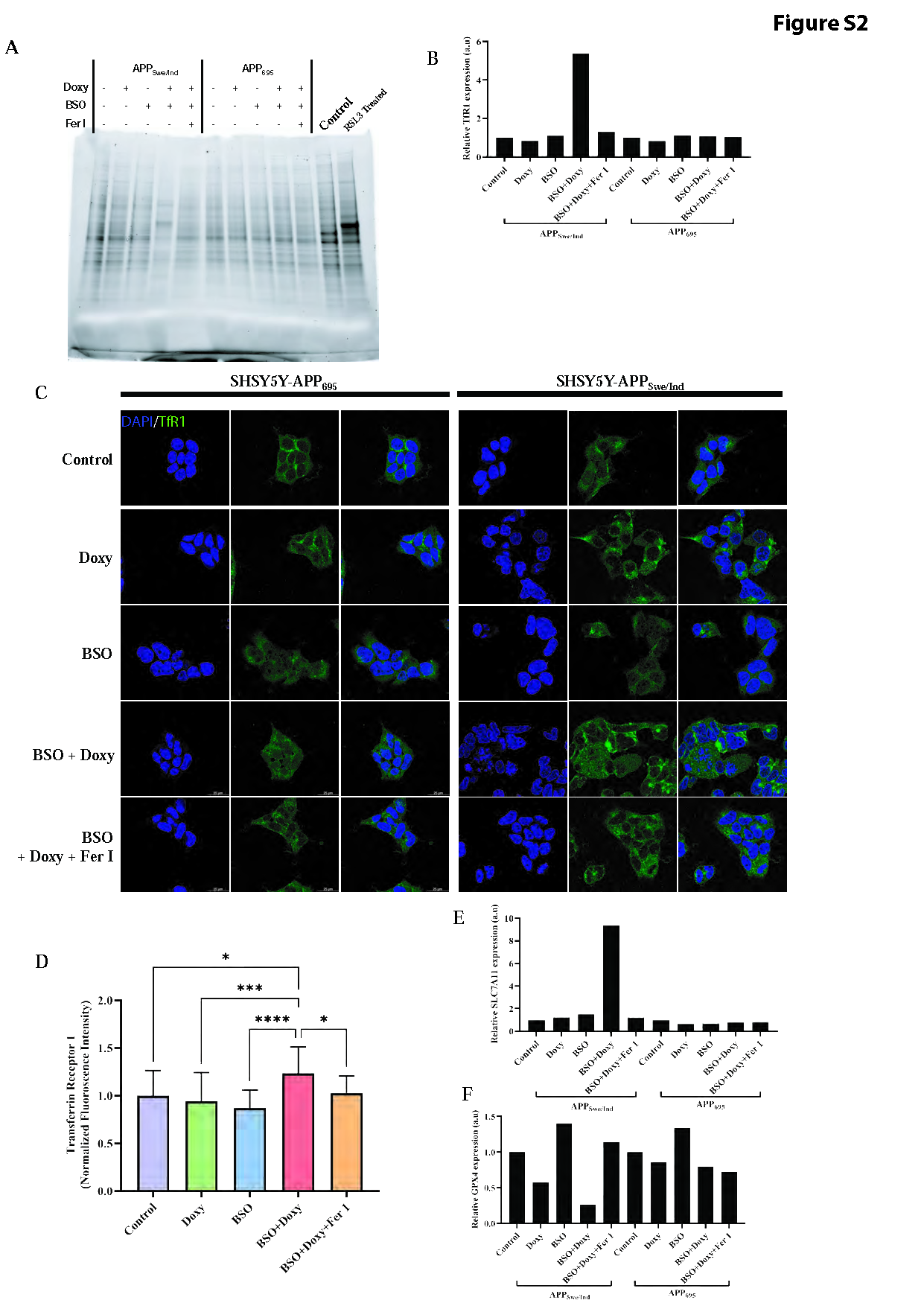

### Supplementary figure 2

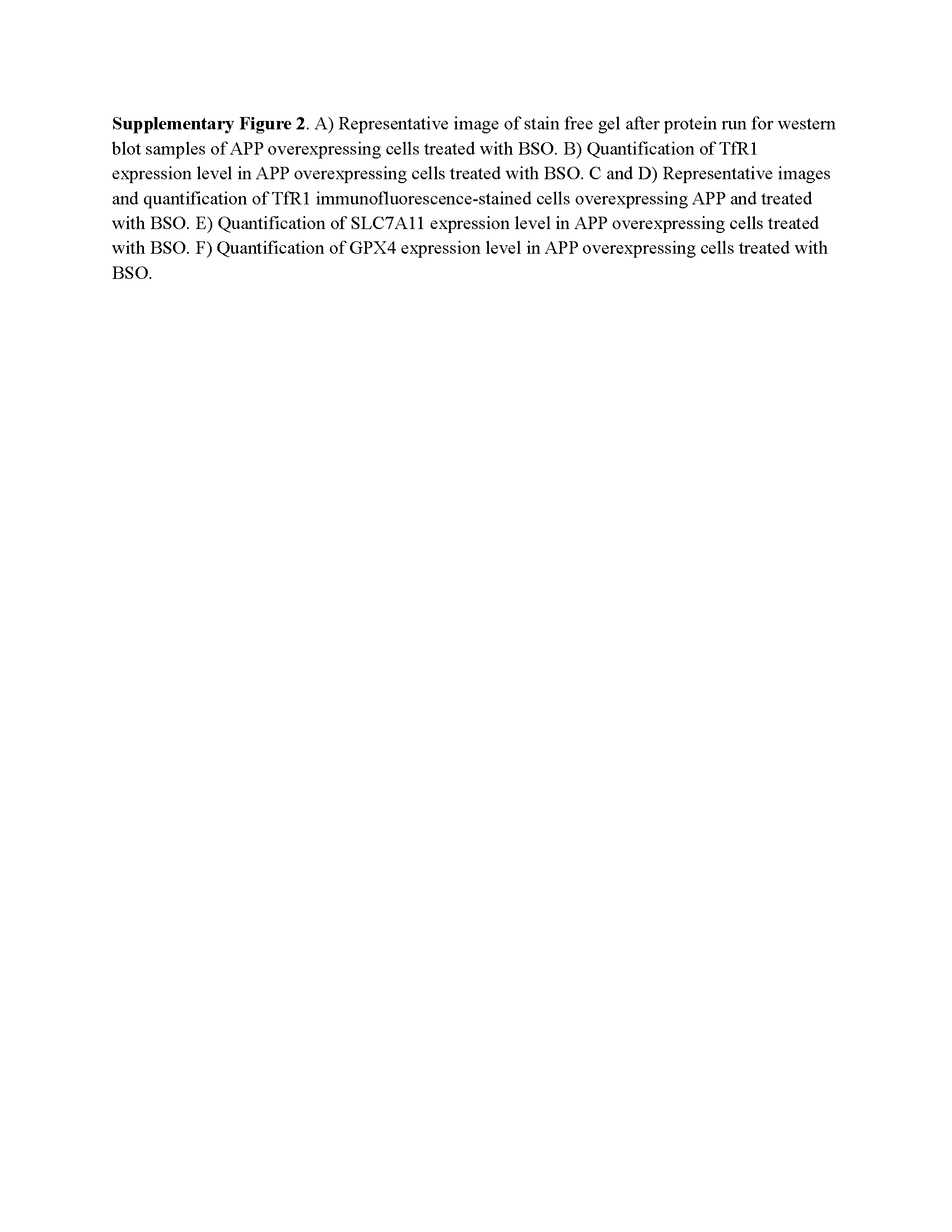

### Supplementary figure 3

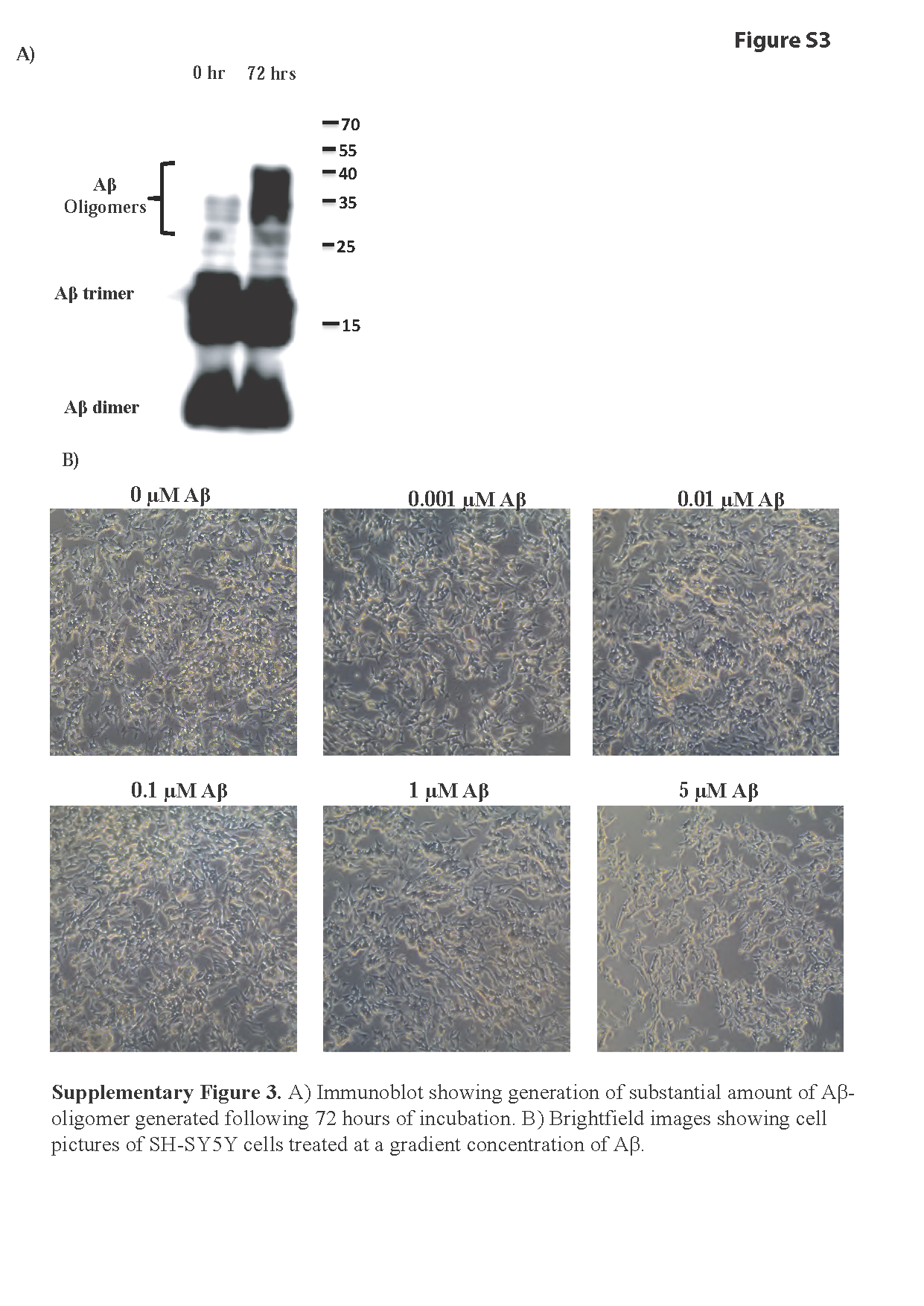

### Supplementary figure 4

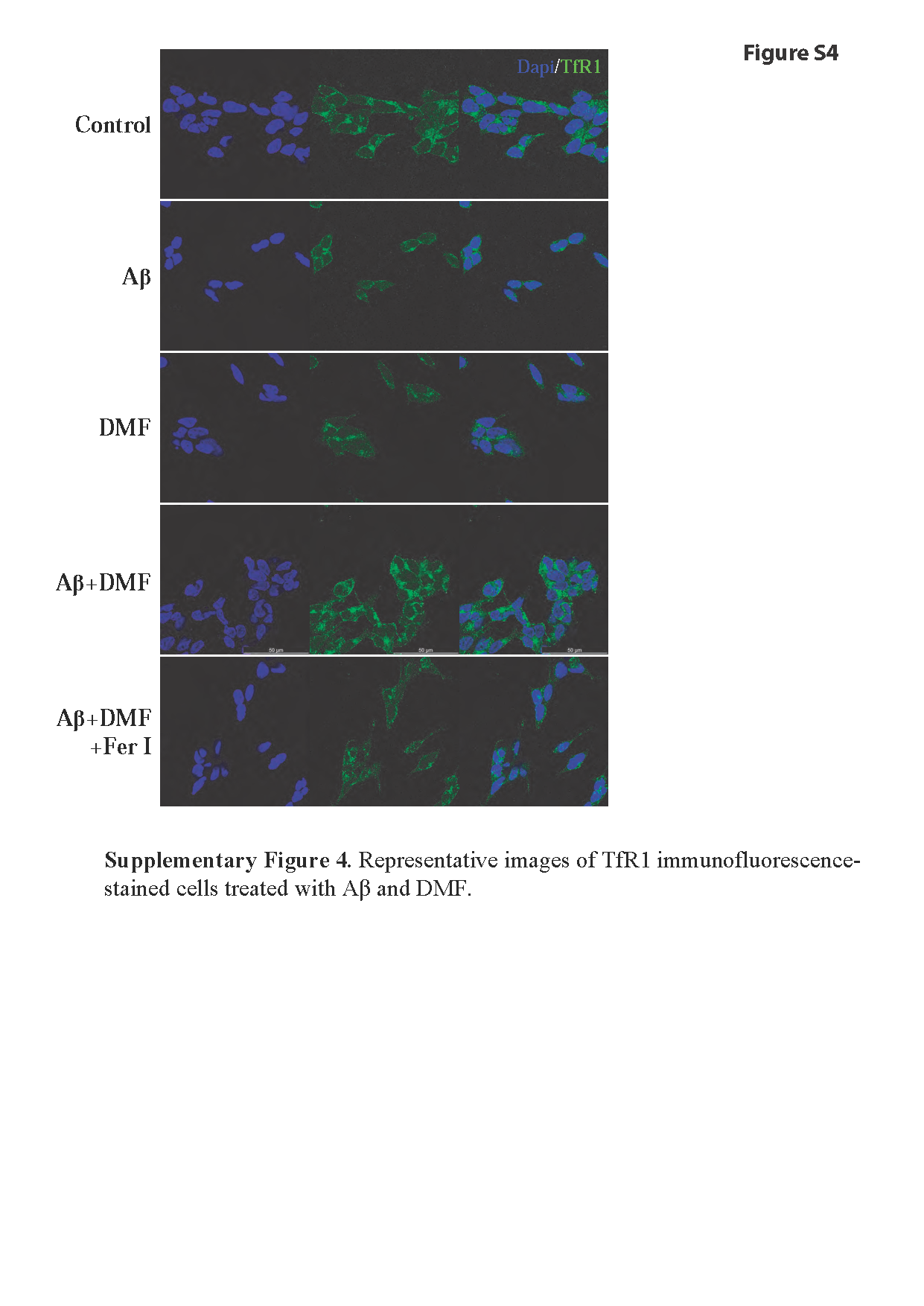

### Supplementary figure 5

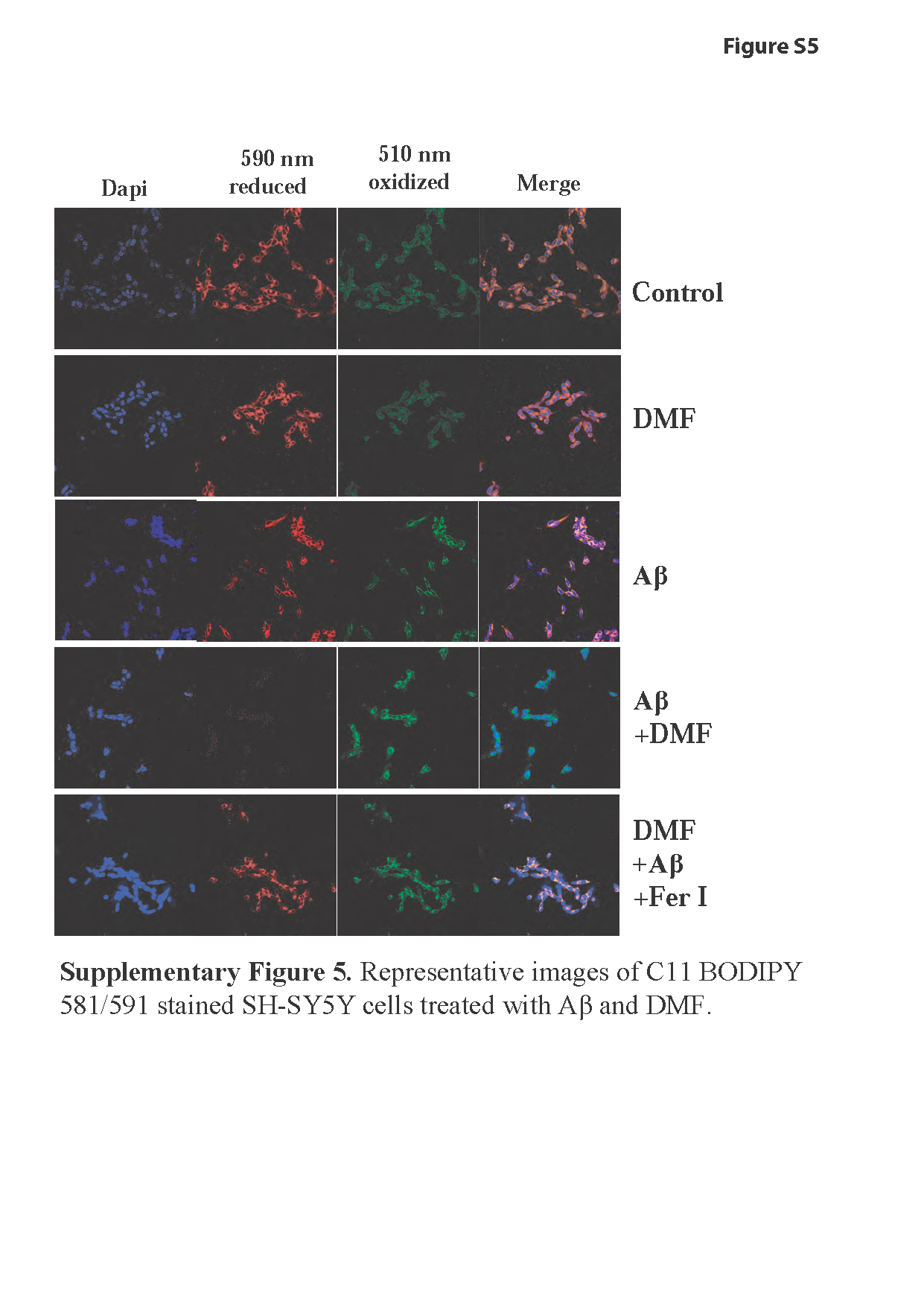

### Supplementary figure 6

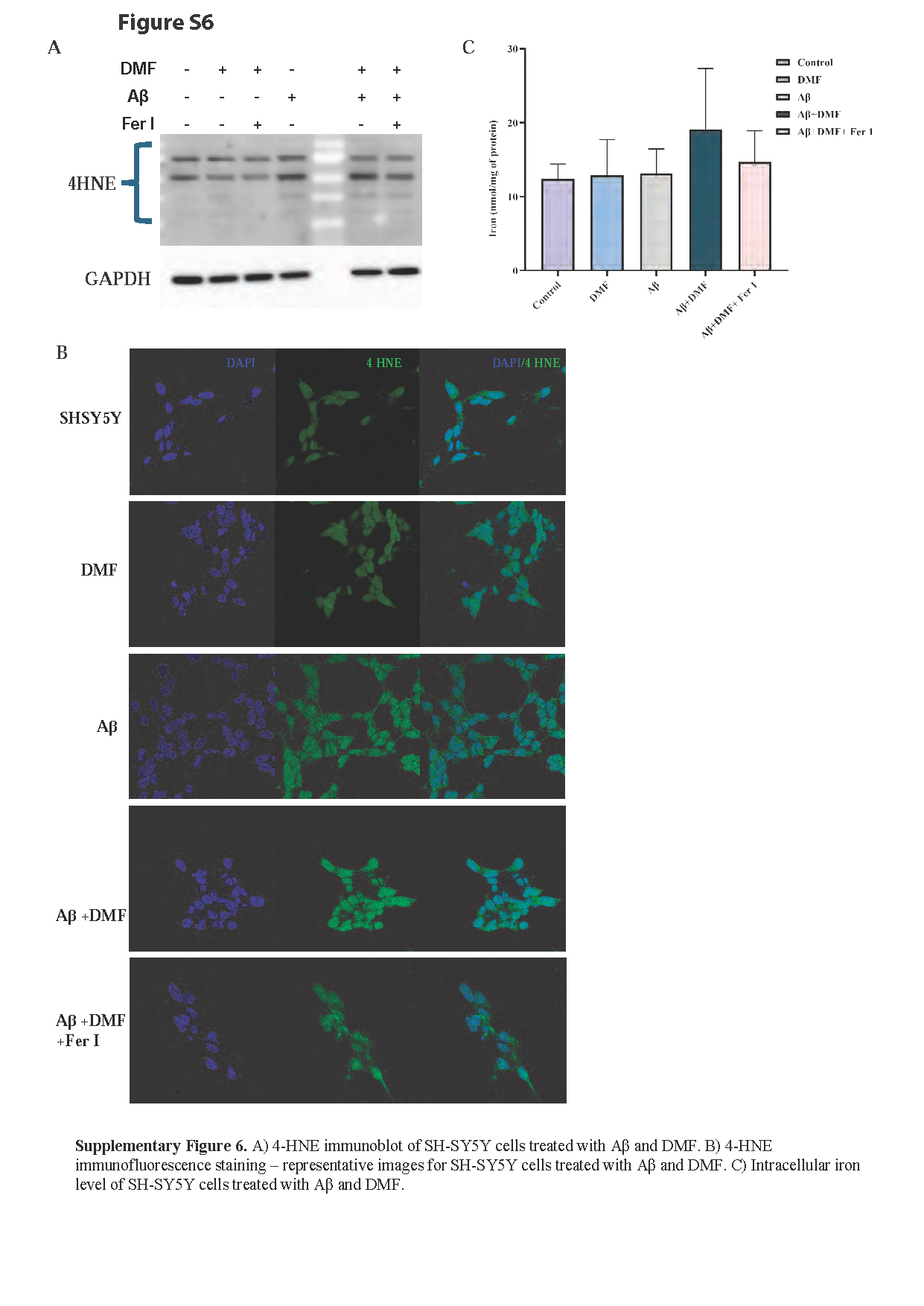

### Supplementary figure 7

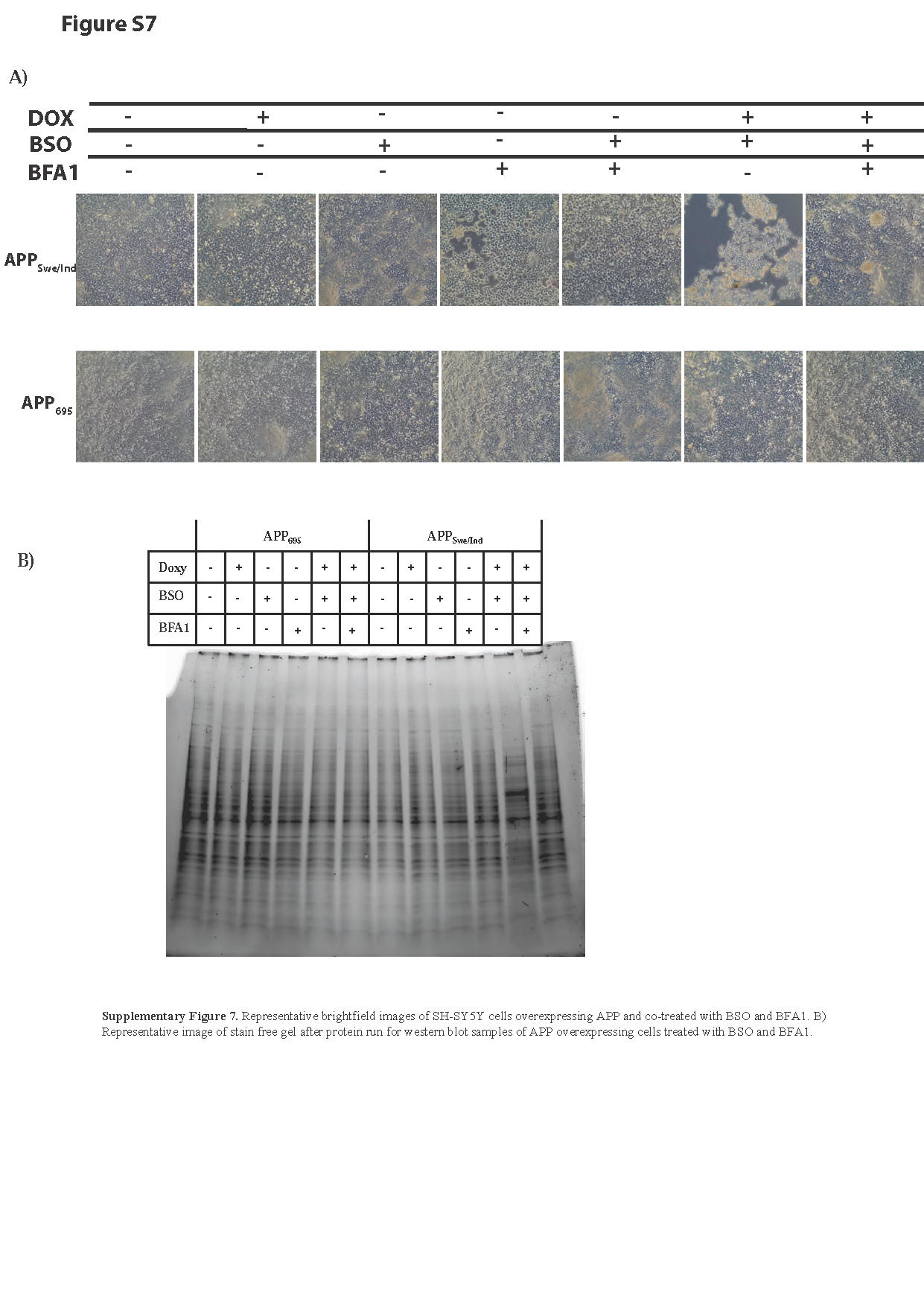
